# Bioengineered human bone-mimetic niche compositions regulate breast cancer cell quiescence and therapy response

**DOI:** 10.64898/2025.12.18.695089

**Authors:** Evrim C. Kabak, Manuele G. Muraro, Sok L. Foo, Ewelina M. Bartoszek, Andres Garcia-Garcia, Atharva Damle, Florian Pouzet, Mohamed Bentires-Alj, Ivan Martin

## Abstract

Disseminated tumor cells (DTCs) in the bone are widely regarded as the cellular seeds of late metastatic relapse in estrogen receptor-positive (ER+) breast cancer (BC). However, how distinct bone compartments influence BC cell quiescence and endocrine therapy response remains unclear. Mechanistic insight has been hindered by limited access to human bone samples and by the lack of relevant human models that permit controlled manipulation of stromal compartments. Here, we developed a fully human, 3D bone-mimetic system for modular, controllable assembly of engineered osteoblastic (eON), vascularized (eVN), and vascularized osteoblastic (eVON) niche compositions. These niches were generated by perfusion culture of human bone marrow-derived mesenchymal stromal cells (hBM-MSCs) and/or human adipose tissue-derived stromal vascular fraction (hAT-SVF) cells within porous ceramic scaffolds. The resulting tissue microenvironments were then used as a substrate for the culture of an ER+ BC cell line, expressing a mutant reporter of p27 to monitor the quiescent status. We found that the eON enhanced BC cell proliferation, whereas the eVN was enriched in quiescent BC cells positive for NR2F1, a dormancy-associated transcription factor, and located near perivascular elements. Treatment with the selective ER degrader fulvestrant reduced BC cell numbers in vascularized niches (eVN and eVON) but not in eON, despite comparable receptor degradation. In summary, we developed a modular human platform for dissecting niche-specific regulation of BC quiescence, proliferation, and endocrine therapy response. The system can be further used to investigate perivascular niche-dependent mechanisms of BC cell dormancy and to guide the development of therapeutic strategies preventing recurrence in ER+ BC patients.

## Introduction

Bone is the predominant site of metastatic relapse in estrogen receptor-positive (ER+) breast cancer (BC) [1-4]. Disseminated tumor cells (DTCs) are frequently detected in bone marrow (BM) aspirates of patients lacking clinically detectable metastases. The presence of DTCs in the bone compartments correlates with poor prognosis, highlighting their potential as seeds for future metastases [5-9]. Understanding the contribution of bone niches to DTC outgrowth is therefore critical for developing strategies to prevent metastatic relapse.

Within the bone, DTCs localize to specialized microenvironments, primarily the osteogenic and perivascular niches [10, 11]. The osteogenic niche comprises osteoblast- and osteoclast-lineage cells lining the bone surface [12]. The osteoclast-driven “vicious cycle” of advanced stage of bone metastasis, marked by tumor-induced osteolysis and release of growth factors that promote further tumor growth, is well-characterized [2, 9, 13-16]. Once the osteolytic lesions emerge, treatment is largely palliative [17-21]. In contrast, the contribution of osteoblasts during early metastatic stages is understudied [12, 22]. Growing evidence suggests that osteoblasts promote early colonization, therapeutic resistance, and proliferation of ER+ BC cells [23-25]. Targeting these early cellular interactions, before the onset of irreversible osteolysis, may represent a more effective therapeutic window.

The other specialized bone compartment to which DTCs mainly traffic is the perivascular niche, which is often anatomically adjacent to or overlapping with the osteogenic niche [26]. While the osteogenic niche is primarily associated with promoting DTC proliferation during early metastatic progression, prior work has shown that the perivascular niche can sustain DTC dormancy [27]. Using intravital imaging on bone marrow (BM) of a BC xenograft model, DTCs were found predominantly localized near perisinusoidal vessels. Analyses of patient BM samples also revealed that non-proliferative, Ki67-negative (Ki67-) DTCs are more frequently found adjacent to sinusoidal vessels than endosteal surface [26]. Thrombospondin 1 (TSP1) expression was strongly associated with BC cell dormancy in an *in vitro* organotypic model composed of endothelial and stromal cells, mimicking the perivascular niche [27]. Despite their distinct features, the osteogenic and perivascular niches are interconnected through shared mesenchymal progenitors that give rise to both endothelial and osteoblastic lineages. Indeed, specialized type H capillaries, enriched in CD31 and endomucin (encoded by *Emcn* in mice), typically reside near the endosteum and contribute to both angiogenesis and osteogenesis via factors such as Noggin [10, 28-30]. These developmental and anatomical overlaps underscore the importance of not only dissecting the individual effects of osteogenic and perivascular microenvironments on DTC outgrowth, but also of modeling transitional zones, where osteogenic and vascular cues converge.

Although recent *in vivo* studies have advanced understanding of how discrete bone compartments influence DTC outgrowth, they also underscore the need for fully human *ex vivo* systems to dissect and compare niche-specific contributions to metastatic progression. A central limitation is the scarcity of solitary DTCs in mouse bone, which hampers spatial and functional interrogation of discrete bone niches during early metastatic stages. Additionally, murine models often fail to recapitulate spontaneous tumor cell dissemination and are further constrained by the rarity of ER+ mammary tumor cell lines compatible with immunocompetent hosts [31-33]. Complementing these challenges, existing *in vitro* and *ex vivo* models often lack the modularity to independently engineer bone-mimetic niches and the dynamic perfusion required for physiological delivery of oxygen, nutrients, and drugs [27, 34-40]. Together, these limitations restrict the ability to resolve niche-specific effects on BC cell dormancy and reactivation, particularly in ER+ disease, where the underlying mechanisms remain poorly understood.

We have previously developed a human, bioreactor-based 3D BM niche model using hydroxyapatite scaffolds seeded with human BM-derived MSCs (hBM-MSCs), and/or adipose tissue-derived stromal vascular fraction (hAT-SVF) cells [41-43]. These engineered constructs mimic the mineralized architecture of trabecular bone with or without vascularization. They also reproduce key features of mineralized and perivascular bone regions [42, 44, 45].

In this study, we investigated the influence of distinct engineered bone-mimetic niches on BC cell quiescence by adapting our previously established 3D perfusion-based culture systems. This platform enables modular engineering of osteogenic, vascular, and vascularized osteogenic bone niches under controlled conditions. We used MCF7-tdTomato/mVenus-p27K^-^ cells to track BC cells in G0 cell cycle arrest by a mutant p27 reporter [46, 47]. To validate niche-specific dormancy, we further evaluated the expression of nuclear receptor subfamily 2, group F member 1 (NR2F1), an orphan nuclear receptor used as a functional dormancy marker in metastatic mouse models and a prognostic biomarker of dormant DTCs in the BM of BC patients [48-50]. Indeed, our side-by-side comparative approach revealed that nuclear NR2F1+ MCF7 cells were enriched in eVN, underscoring its utility as a platform to study cell-extrinsic regulation of BC dormancy in bone.

## Results

### Engineered osteoblastic niche promotes MCF7 cell proliferation

To assess the effects of an osteogenic niche on BC cell quiescence and proliferation, hBM-MSCs were seeded on hydroxyapatite scaffolds and expanded for one week, followed by three weeks of osteogenic differentiation to engineer an osteoblastic niche (eON) [41, 43, 44]. MCF7-tdTomato/mVenus-p27K^-^ cells were then seeded and cultured for two weeks under perfusion in the eON or in the niche-free scaffold (NFS) as control (Fig. 1A) [46, 47]. The total number of MCF7-tdTomato/mVenus-p27K^-^ cells increased after two weeks of culture in the eON compared to NFS (Fig. 1B). Immunofluorescence (IF) staining for tdTomato and mVenus further revealed a reduced fraction of quiescent (mVenus+) MCF7 cells in eON (Fig. 1C). Consistently, Ki67 staining showed a higher proportion of proliferating MCF7 cells in eON compared to the control, highlighting that eON enhances MCF7 cell proliferation (Fig. 1D). The eON was previously validated by matrix deposition of collagen type I alpha 1 (COL1A1) and osteocalcin (OCN) [44, 45].

**Figure 1.**
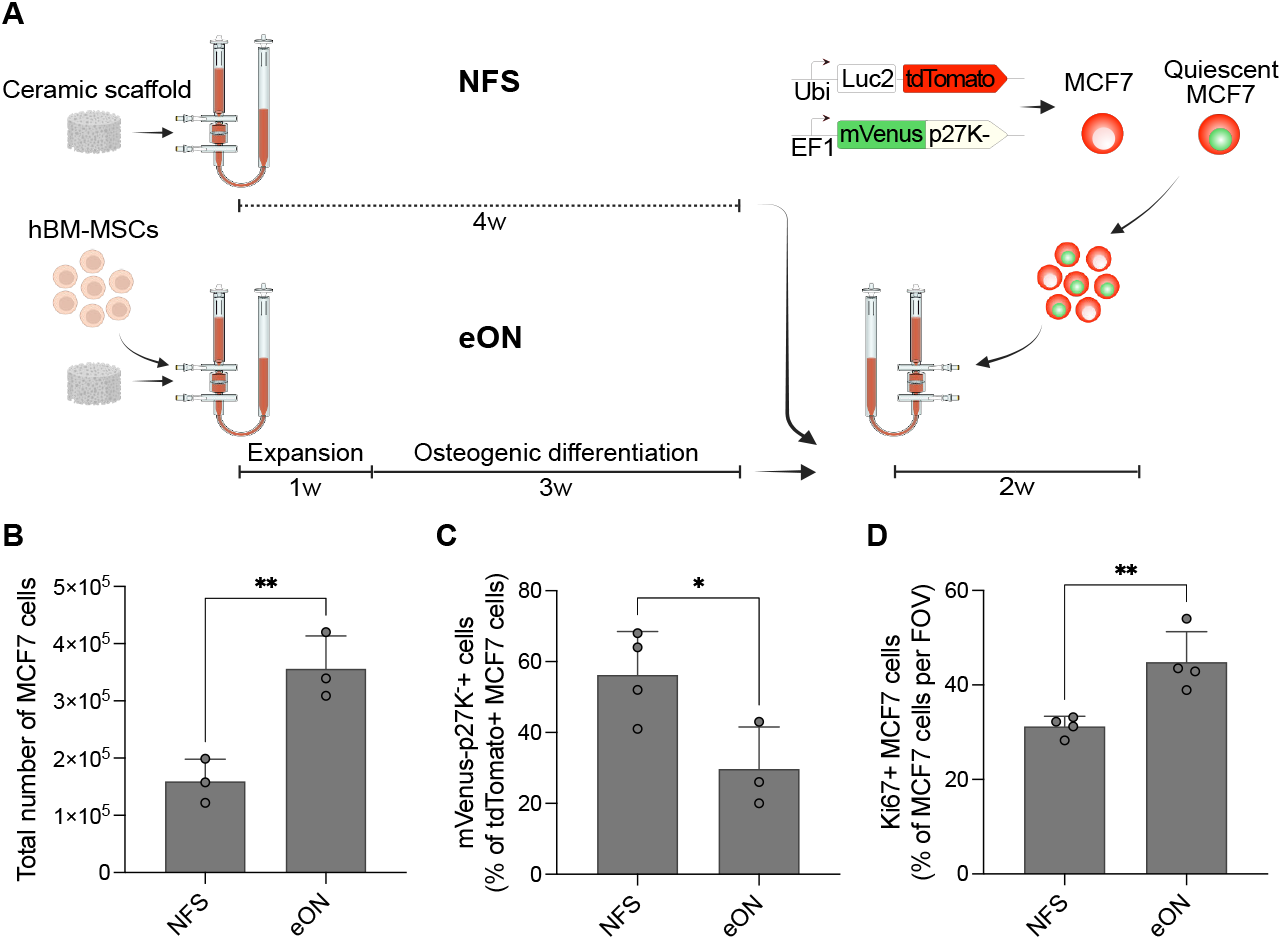
Engineered osteoblastic niche (eON) promotes MCF7 cell proliferation. **(A)** Experimental scheme to generate eON co-cultured with MCF7-tdTomato/mVenus-p27K^−^ for two weeks in perfusion bioreactors. **(B)** Flow cytometry quantification of MCF7 cells after two weeks of culture in niche-free scaffold (NFS) or eON. **(C, D)** Percentage of **(C)** quiescent (mVenus-p27K^−^) MCF7 cells, and **(D)** proliferating (Ki67+) MCF7 cells relative to total tdTomato+ MCF7 cells per field of view (FOV) in NFS or eON. Data presented as mean ± SD, n=3. Statistical significance was determined by two-tailed, unpaired, parametric t test. *p<0.05, **p<0.01. **(C, D)** 5 FOV analyzed per sample.

COL1A1 and OCN staining confirmed osteogenic differentiation in eON, whereas NFS lacked detectable levels of these markers (*SI Appendix*, Fig. *S1*). These experiments validated the feasibility of culturing MCF7 cells in an engineered niche as a prerequisite to evaluate how bone niche compositions influence BC cell cycle states.

### eON, eVN, and eVON model distinct osteoblastic and vascular features of bone

Although prior studies have suggested that the perivascular niche of the bone may contribute to BC dormancy, the specific influence of vascular and perivascular cues on BC cell cycle state, particularly in the presence of osteogenic elements, has not yet been addressed [26, 27, 51]. To address this, we engineered two additional environments: a vascularized niche (eVN) and a vascularized osteogenic niche (eVON). eVN was generated by seeding hAT-SVF cells into hydroxyapatite scaffolds and culturing them for two weeks in medium supplemented with fibroblast growth factor 2 (FGF2). For eVON, hAT-SVF cells were seeded into a pre-osteogenic niche and cultured in osteogenic medium supplemented with FGF2 for two weeks [42]. Perfusion bioreactors ensured uniform distribution of MCF7-tdTomato/mVenus-p27K^-^ cells upon subsequent seeding. MCF7 cells were then introduced into each niche configuration and co-cultured for an additional two weeks (Fig. 2A).

**Figure 2.**
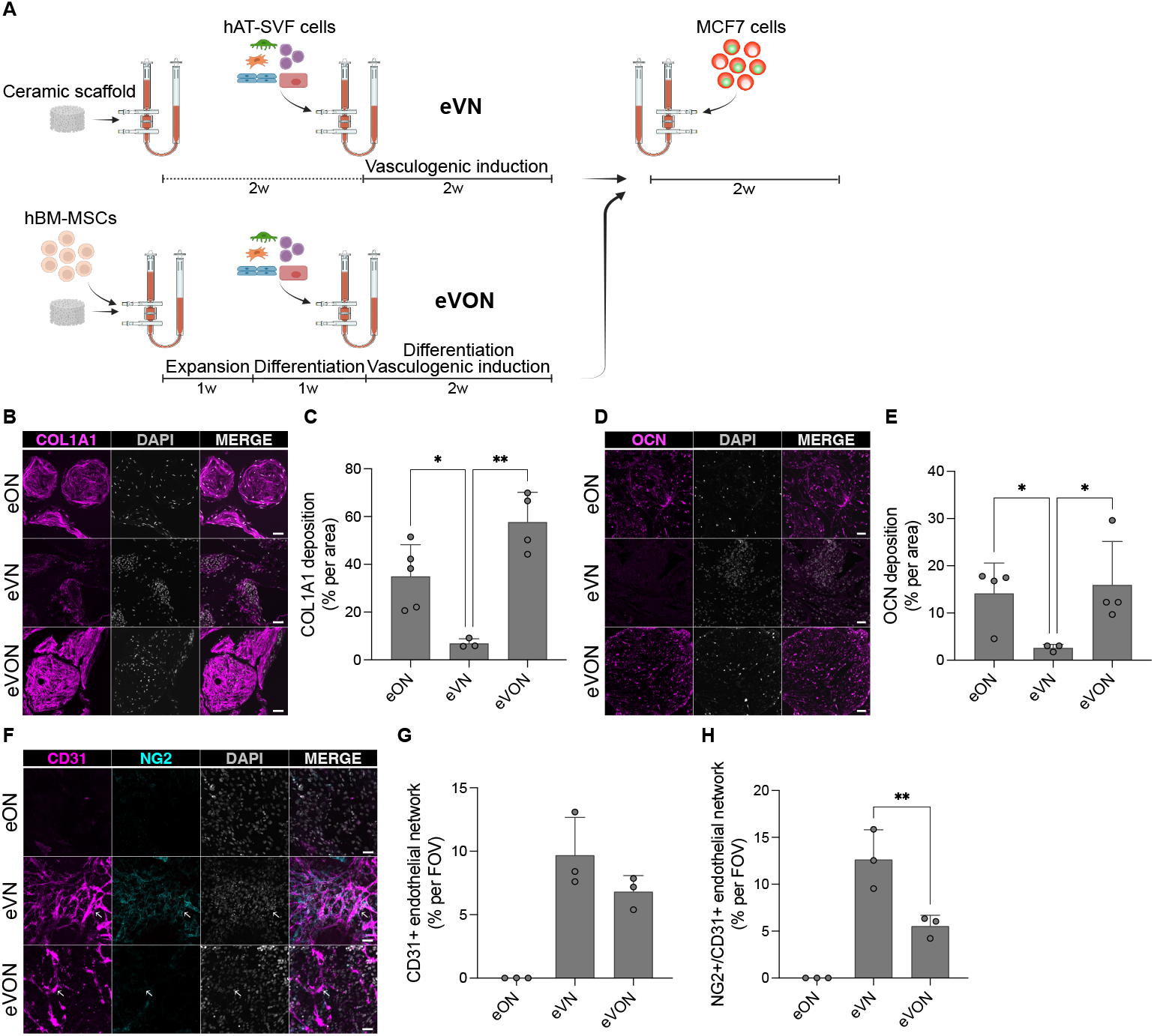
eON, eVN and eVON recapitulate osteoblastic, vascularized, and vascularized osteoblastic bone microenvironments, respectively. **(A)** Experimental scheme illustrating generation of eVN and eVON co-cultured with MCF7-tdTomato/mVenus-p27K^−^ cells for two weeks. **(B)** Representative IF staining and **(C)** quantification of COL1A1 deposition (percentage per area) across the niches. **(D)** Representative IF staining and **(E)** quantification of OCN deposition (percentage per area) across the niches. Magnification: 20x; scale bar: 50 µm. **(F)** Representative whole-mount immunofluorescence (IF) staining of CD31 (magenta) and NG2 (cyan) in eON, eVN, and eVON. Magnification: 20x; scale bar: 50 µm. **(G, H)** Quantification of **(G)** CD31+ endothelial network density (percentage per FOV) and **(H)** NG2+ perivascular cell density on CD31+ endothelial network **(H)** (percentage per FOV) in eON, eVN, and eVON. **(C, E, G, H)** Data presented as mean ± SD, n=3-5 (two independent experiments). **(G, H)** 5 FOV analyzed per sample. Statistical significance was determined by one-way ANOVA followed by Tukey’s multiple comparison test. *p<0.05, **p<0.01. Multiple comparisons were performed exclusively between eVN and eVON for vascular analysis.

To determine whether the engineered niches recapitulate key features of bone microenvironments, we assessed osteogenic matrix deposition (i.e., COL1A1 and OCN) and endothelial (i.e., CD31) network formation. The osteogenic compartment of the three niches was characterized by IF staining and quantified using an optimized image analysis pipeline (*SI Appendix* Fig. *S2A, S2B*). The highest levels of COL1A1 deposition were observed in eVON (57.8 ± 12.4%), followed by eON (35.0 ± 13.3%), and lowest in eVN (6.9 ± 1.9%) (Fig. 2B, 2C), likely due to the higher stromal cell content in eVON. OCN deposition was low in eVN (2.6 ± 0.8%), but higher in both eON (14.1 ± 6.4%) and eVON (16.0 ± 9.2%) (Fig. 2D, 2E). These results confirm that eON and eVON retain osteogenic matrix features compared to eVN.

Staining for CD31 revealed the self-organized, branched endothelial networks in both eVN and eVON after two weeks of co-culture with MCF7-tdTomato/mVenus-p27K^-^ cells (Fig. 2F). Notably, these networks formed and persisted without exogeneous angiogenic factor supplementation, indicating stable endothelial organization. eON lacked CD31+ endothelial structures, consistent with the avascular nature of mineralized osteoblastic niches (Fig. 2F). Using the pericyte marker neuron-glial antigen 2-positive (NG2+; also known as chondroitin sulfate proteoglycan 4, CSPG4), we measured proportion of perivascular cells associated with CD31+ structures using an optimized image analysis pipeline (*SI Appendix* Fig. *S2C*). CD31+ network density was comparable between eVN and eVON (Fig. 2G). However, NG2+ cells more frequently colocalized with CD31+ networks in eVN, reflecting a more developed perivascular architecture (Fig. 2F, 2H). These findings indicate that endothelial network-forming capacity is maintained in both niches, while perivascular organization is less extensive in eVON.

### Nuclear NR2F1+ MCF7 cells are enriched in eVN

Next, we assessed the effect of distinct cellular compositions provided by eON, eVN, and eVON on MCF7-tdTomato/mVenus-p27K^-^ cell quiescence and proliferation. To this end, we first quantified the number of MCF7 cells retrieved from each niche after two weeks of co-culture. Flow cytometry analysis revealed the highest number of MCF7 cells in eVON, followed by eON, with eVN yielding the lowest cell numbers. These findings suggest that combined osteoblastic and vascular components correlate with enhanced MCF7 cell count (Fig. 3A).

**Figure 3.**
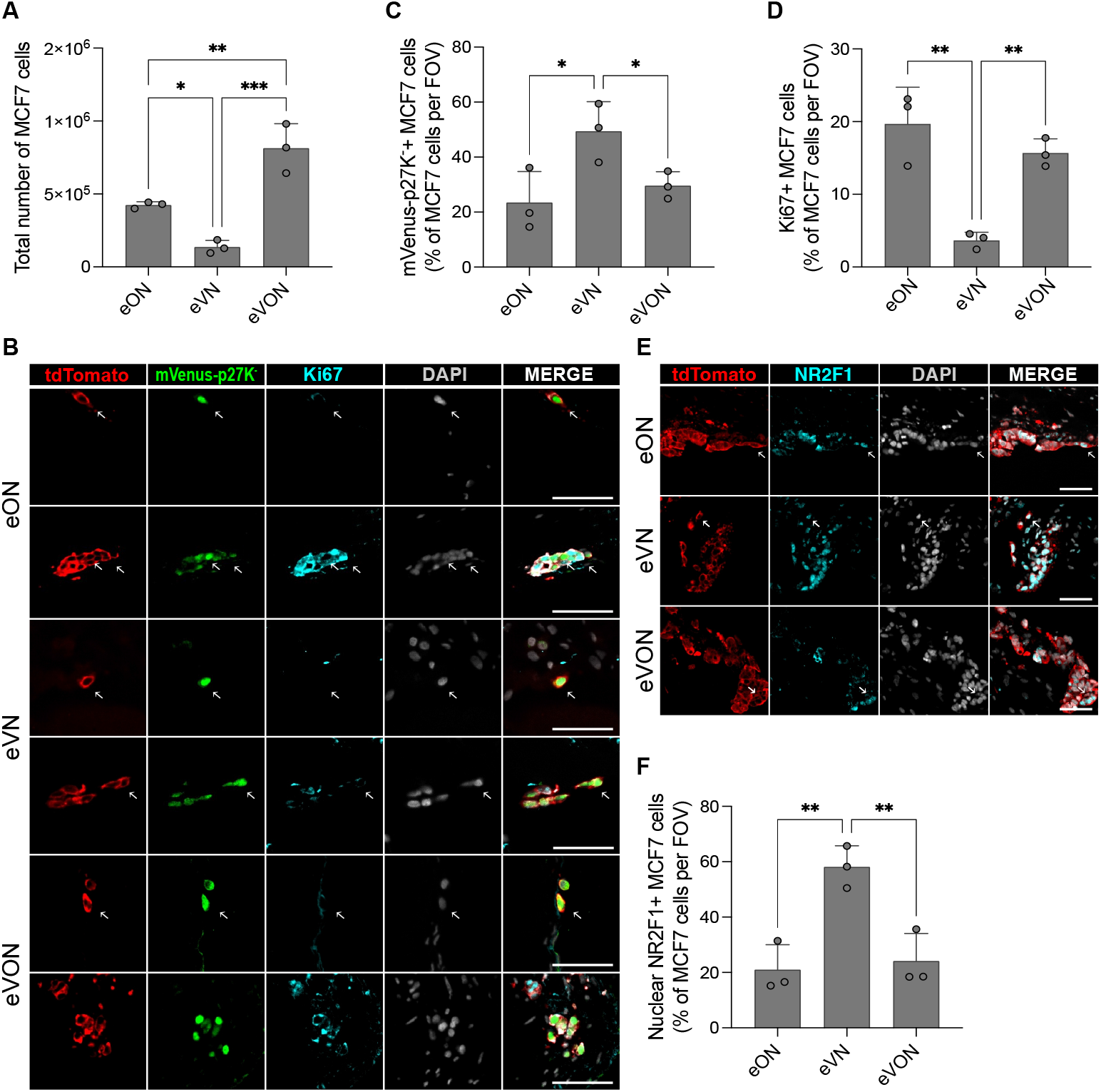
eVN enriches for quiescent, non-proliferative MCF7 cells with elevated nuclear NR2F1. **(A)** Flow cytometry quantification of MCF7-tdTomato/mVenus-p27K^-^ cells after two weeks of co-culture with eON, eVN and eVON. **(B)** Representative IF staining showing quiescent (mVenus-p27K^−^+; green) MCF7 cells appearing as solitary and clustered with proliferating (Ki67+; cyan) MCF7 cells (tdTomato+; red) in eON, eVN and eVON after two weeks of co-culture. **(C, D)** Percentage of **(C)** quiescent (mVenus-p27K^-^+) MCF7 cells, and **(D)** proliferating (Ki67+) MCF7 cells relative to total MCF7 cells per FOV. **(E)** Representative IF staining of NR2F1 (cyan) and for tdTomato (red) after two weeks of co-culture in eON, eVN, and eVON. Magnification: 20x; scale bar: 50 µm. **(F)** Percentage of nuclear NR2F1+ MCF7 cells relative to total MCF7 cells per FOV in the respective niches. **(A, C, D, F)** Data presented as mean ± SD, n=3 (three independent experiments; each data point represents the mean of three technical replicates per donor). Statistical significance was determined by one-way ANOVA with Tukey’s multiple comparison test. *p<0.05, **p<0.01, ***p<0.001. **(C, D, F)** 5 FOV analyzed per sample.

IF staining showed that mVenus-p27K- and Ki67 expression were mutually exclusive, confirming the specificity of the quiescence reporter (mVenus-p27K^-^) (Fig. 3B). Both quiescent (tdTomato+/mVenus+/Ki67-) and proliferating (tdTomato+/mVenus-/Ki67+) cancer cells were detected across all niches. Quiescent cancer cells were also found as solitary cells or as clusters with proliferative cells in all niches (Fig. 3B, *SI Appendix*, Fig. *S3*). Quiescent and proliferating cancer cells were quantified using an optimized image analysis pipeline (*SI Appendix*, Fig. *S4*). eVN had the highest proportion of quiescent MCF7 cells relative to eON and eVON (Fig. 3C). Coherently, it had the lowest fraction Ki67+ MCF7 cells (Fig. 3D).

Given the higher percentage of quiescent MCF7-tdTomato/mVenus-p27K^-^ cells in eVN, we asked whether this phenotype was associated with a dormancy-linked transcriptional program. Prior studies have shown that microenvironmental cues in the bone, including cytokine-mediated activation of p38 mitogen-activated protein kinase (MAPK) signaling, can induce dormancy through upregulation of specific transcriptional regulators such as NR2F1 [48-50, 52]. Notably, we found nuclear colocalization of mVenus-p27K^-^ and NR2F1 expression (*SI Appendix*, Fig. *S5*) and more frequently within the eVN compared to eON and eVON (Fig. 3E, F), suggesting that eVN promotes dormancy phenotype. Furthermore, the higher abundance of NG2+/CD31+ endothelial network in eVN correlated with increased frequencies of nuclear NR2F1+ and quiescent MCF7 cells.

### Fulvestrant-induced MCF7 cell number reduction is confined to eVN and eVON

To test whether distinct niche compositions influence therapeutic response, we treated engineered co-cultures with fulvestrant for one week. Qualitative assessment of ERα Immunohistochemistry (IHC) revealed a pronounced loss in nuclear ERα staining in MCF7 cells across all three conditions (Fig. 4A). Consistent with this, the percentage of nuclear ERα+ MCF7-tdTomato/mVenus-p27K^-^ cells was markedly reduced across all three niches, confirming target engagement (Fig. 4B). Notably, MCF7 cell numbers significantly decreased in eVN and eVON upon treatment, but remained unchanged in eON (Fig. 4C), indicating that fulvestrant-induced reduction in cell number was confined to the niches incorporating endothelial networks.

**Figure 4.**
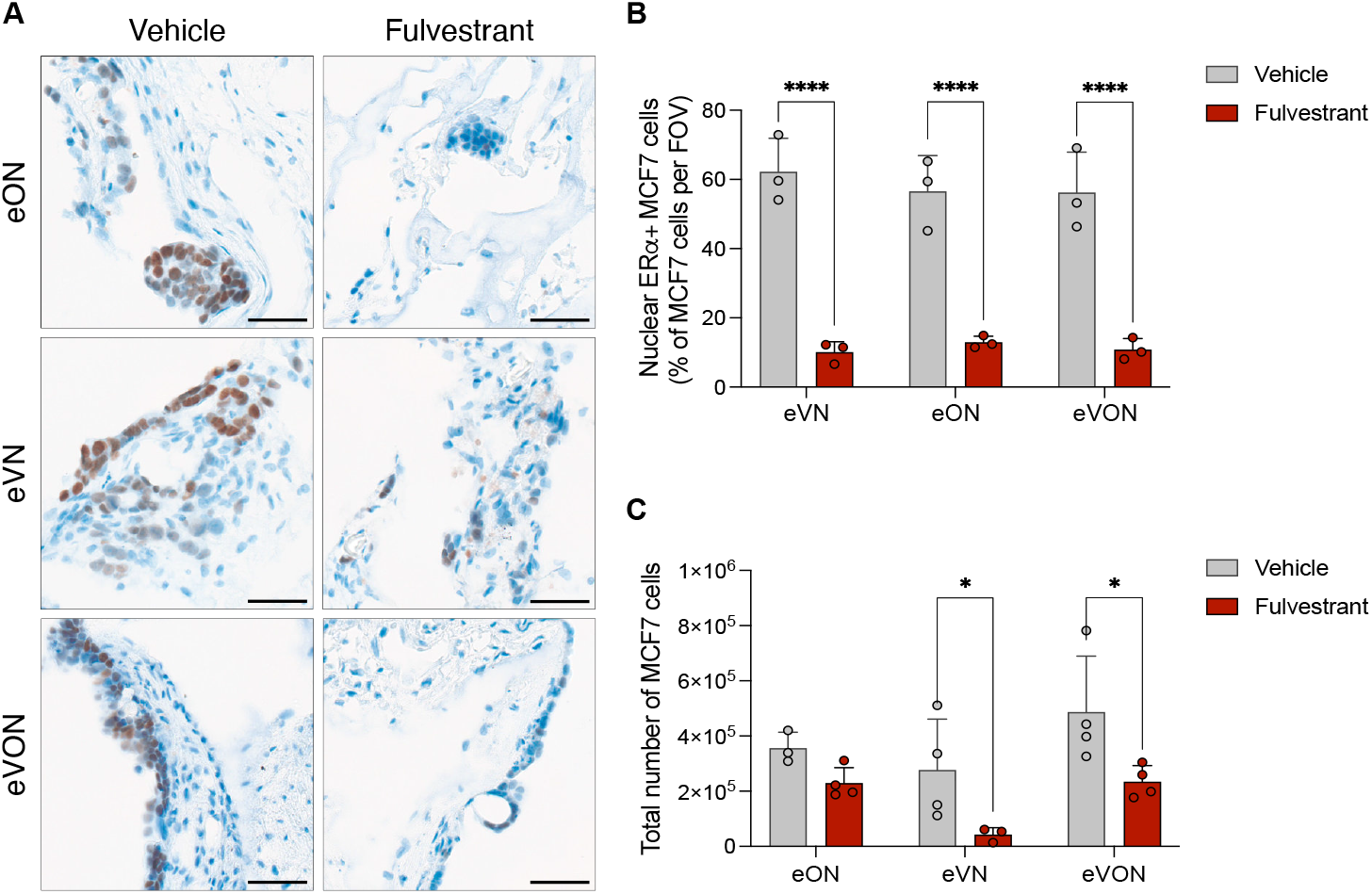
Fulvestrant reduces MCF7 cell numbers only in vascularized niches, despite comparable ERα degradation across all conditions. **(A)** Representative IHC staining images of ERα and **(B)** percentage of nuclear ERα+ MCF7-tdTomato/mVenus-p27K-cells relative to total MCF7 cells per FOV upon one week of treatment in eON, eVN and eVON. **(C)** Flow cytometry quantification of MCF7 cells after one week of treatment in eON, eVN and eVON. Data presented as mean ± SD, n=3-4 (two independent experiments). Magnification: 40x; scale bar: 50 µm. **(A, B)** Statistical significance was determined by two-way ANOVA with Tukey’s multiple comparison test. *p<0.05, ****p<0.0001.

## Discussion

In this study, we developed a fully human, modular 3D bone-mimetic model to investigate cancer-stroma interactions during BC bone metastatic progression. This platform allows the stepwise introduction of microenvironmental complexity and supports direct comparison of different cellular components in a modular fashion under controlled conditions. Importantly, our model allows cancer cells to be seeded after bone-mimetic niche formation thanks to perfusion flow, thereby better mimicking the temporal sequence of spontaneous cancer cell dissemination. This design enables spatial analysis of niche-specific effects on cancer cell cycle states, including quiescence and proliferation, in a physiologically relevant and experimentally tractable context.

Our findings that the eON promotes MCF7-tdTomato/mVenus-p27K^-^ cell proliferation are consistent with previous *in vivo* studies [24] showing that osteoblast-rich microenvironments enhance early colonization and proliferation of BC cells in the bone [23, 24]. This underscores the often-overlooked contribution of osteoblasts to the early, non-osteolytic stages of metastatic progression, and highlights the importance of modeling these niches independently of the osteoclast-driven vicious cycle.

The eVN and eVON enabled us to investigate the influence of endothelial and perivascular elements on cancer cell quiescence in the absence or presence of an osteoblastic niche. eVN yielded the lowest total number of MCF7 cells yet contained a higher fraction of quiescent cells than eON and eVON. In contrast, eVON supported the highest MCF7 cell numbers, while the quiescent fraction was reduced relative to eVN. This pattern suggests that osteogenic cues may modulate the impact of vascular elements on BC cell quiescence, potentially by interfering with perivascular organization [10, 25, 28].

Prior studies have shown that specialized capillaries in the BM, such as type H vessels coordinate osteogenesis through endothelial Notch signaling [28, 29]. Disruption of these vascular-osteogenic interactions may alter the microenvironmental signaling, potentially affecting DTC outgrowth. Importantly, the NG2+/CD31+ networks resemble the dormancy-supportive vasculature, where DTCs adjacent to stable microvessels had elevated p27 expression and remained quiescent, in contrast to reactivation near sprouting vessels [27]. Perivascular cells expressing NG2 were more frequently associated with CD31+ endothelial networks in eVN than in eVON, suggesting a more stable engineered vasculature in the absence of osteogenic differentiation. Whether this contributes to the higher proportion of quiescent cells in eVN compared to eVON remains to be addressed.

In line with the enrichment of the quiescent cell fraction in eVN, we also observed a significantly higher proportion of nuclear NR2F1+ cells in this niche compared to eON and eVON. BM-derived soluble factors, including TGF-β2 and BMP7, particularly enriched in perivascular regions, can activate the p38 MAPK pathway and upregulate dormancy-associated genes such as NR2F1 [48, 49, 52-54]. This association suggests that structural features of the eVN niche, including enriched NG2+/CD31+ networks, may potentially contribute to the activation of dormancy-related signaling programs. Together, these results validate the pathophysiological relevance of the eVN and underscore its utility for modeling dormancy-permissive microenvironments.

A striking observation was the presence of quiescent cells both as solitary cells and within clusters that also contained proliferative cells. This spatial distribution mirrors features of both cellular dormancy and tumor mass dormancy [55]. While tumor mass dormancy is defined by a balance between proliferation and apoptosis that limits net growth, we did not assess apoptosis in this study. Thus, we cannot determine whether a dormancy equilibrium exists in the observed clusters. Future studies incorporating apoptotic markers (e.g., cleaved PARP, Annexin V/PI, or TUNEL) and longitudinal live-cell imaging will be necessary to distinguish between cellular dormancy and tumor mass dormancy.

Beyond cell cycle regulation, our system also demonstrates utility for therapeutic testing. Fulvestrant treatment reduced BC cell numbers in eVN and eVON, compared to eON, despite the confirmed nuclear ERα degradation. One potential explanation for the limited response in eON could be fulvestrant-induced changes in ECM, as previous studies have shown that collagen-rich microenvironments can reduce the efficacy of ER-targeted therapies [56, 57]. These data underscore the potential of our platform in dissecting microenvironment-dependent treatment responses and in refining specific therapeutic strategies.

There are limitations specifically regarding the further validation of the dormancy model. Although our data support the presence of NR2F1-associated dormancy in eVN, several important criteria remain unmet for full dormancy validation [58]. Most critically, dormancy is defined not only by G0 cell cycle arrest, but also by its reversibility [59]. Although we observed enrichment of mVenus-p27K^-^+ and Ki67-proportion of MCF7 cells, additional assays are needed to demonstrate the capacity of these cells to re-enter the cell cycle upon niche alteration or exogenous stimulation. Furthermore, the possibility that some quiescent cells are senescent rather than dormant populations must be considered, and should be addressed through assessment of senescence markers such as senescence associated β-galactosidase (SA-β-Gal) or p16INK4a [55, 60].

Another critical gap is the lack of functional perturbation studies to establish causality between niche components and dormancy induction. Although our data suggest that stable perivascular architecture in eVN is associated with NR2F1+ quiescence, functional validation such as selective depletion of endothelial or perivascular cell populations, or genetic manipulation of dormancy-related signaling pathways will be essential to define causality.

Despite these limitations, our system offers a modular platform that recapitulates key phenotypic features of the human bone metastatic microenvironment. It enables direct comparison of distinct niche types and allows for phenotypic interrogation of cancer cell states in a pathophysiologically relevant context. Given its human origin and architectural fidelity, this platform holds promise for in-depth studies on the cellular and molecular determinants of tumor dormancy and for drug testing applications. In particular, it could be used to assess the efficacy of agents targeting dormant cells or preventing metastatic outgrowth in niche-specific contexts, offering a valuable preclinical model for therapeutic development.

## Materials and Methods

### hBM-MSC isolation and culture

Human bone marrow derived mesenchymal stromal cells (hBM-MSCs) were isolated as previously described [41, 44, 45]. Cells were cultured in complete medium (CM) composed of a α-minimum essential medium (αMEM) (Gibco; cat# 22571-020), supplemented with 10% fetal bovine serum (FBS, Invitrogen; cat# 548-62-9), 1% HEPES (1M, Gibco; cat# 15630-056), 1% sodium pyruvate (100 mM), 1% GlutaMAX (100X, Gibco; cat# 35050-061), and 1% Penicillin–Streptomycin (PS, Gibco; cat# 15140-122). Nucleated cells were plated at a density of 5 x 10^3^ cells/cm^2^ and maintained at 37°C in a water-jacketed incubator with 5% CO_2_. CM was supplemented with 5 ng/ml of fibroblast growth factor-2 (FGF-2). Medium was changed twice weekly. hBM-MSCs were selected on adherence and proliferation after one week.

### hAT-SVF cell isolation and culture

Adipose tissue was obtained from three healthy, female donors via liposuction or excision at the University Hospital Basel. Isolation of human adipose tissue derived stromal vascular fraction (hAT-SVF) cells was conducted by enzymatic digestion with collagenase type II (Worthington; cat# LS004176), and followed by several centrifugation and purification steps as described [42, 43, 45]. Cell counting was performed using Acridine Orange/Propidium Iodide Stain (Logos biosystems; cat# F23001) at 1:100 dilution in the cell suspension using Luna-FX7™ Automated Cell Counter (Logos biosystems).

## Ethics statement

Human BM aspirates and adipose tissue were collected with informed consent from healthy donors at the University Hospital Basel. The study was approved by the local ethics committee (Ethikkommission Nordwest-und Zentralschweiz, ref. 78/07) and conducted in accordance with EU ethical guidelines.

### Cancer cell culture

MCF7 cells (ATCC) were cultured in Dulbecco’s Modified Eagle’s Medium (DMEM) high glucose (Sigma; cat# D6429), supplemented with 10% FBS, 1% PS, and 1µg/mL insulin at 37°C with 5% CO_2_. Cell line identity was confirmed by short tandem repeat (STR) sequencing, and routinely tested for mycoplasma contamination.

### Generation of lentivirus and transduced MCF7 cells

Lentiviral particles were generated by co-transfecting HEK293T cells with 2µg each of VSVG envelope plasmid, 2 µg each of third-generation packaging plasmids, and 4µg plasmid DN (pFU-Luc2-tdTomato or pCDH-EF1-mVenus-p27K−), using FuGENE HD (Promega) at a 3:1 (reagent:μg DNA) ratio. Viral supernatants were collected at 48- and 72-hours post-transfection, pooled, filtered, and concentrated using Lenti-X Concentrator (Takara). MCF7 cells were sequentially transduced; first with pFU-Luc2-tdTomato lentivirus and Fluorescence-activated Cell Sorting (FACS)-sorted for tdTomato+ expression (BD Aria), followed by transduction with pCDH-EF1-mVenus-p27K^−^ and puromycin selection 0.75 μg/mL.

### Generation of engineered niches

eON was generated by using hBM-MSC as previously described [41, 44]. Briefly, 0.75×10^6^ hBM-MSCs were seeded into hydroxyapatite scaffolds (Engipore®, Finceramica-Faenza; 4 mm x 8 mm) embedded in perfusion bioreactors. Cells were perfused at 3 mL/min superficial velocity for 24 hours (seeding phase), followed by 0.3 mL/min (culture phase). Constructs were cultured in proliferative medium (PM) consisting of CM supplemented with 100 nM dexamethasone (Sigma; cat# D4902), 0.1 mM ascorbic acid-2-phosphate (Sigma; cat# A92902) and 5 ng/mL FGF-2, followed by three weeks in osteogenic medium (OM) consisting in CM supplemented with 10 nM dexamethasone, 10 mM β-glycerophosphate, and 0.1 mM ascorbic acid-2-phosphate. The medium was changed twice weekly.

eVN was generated by seeding 1×10^5^ hAT-SVF cells into the hydroxyapatite scaffolds embedded in perfusion bioreactors. Cells were exposed to 3 mL/min superficial velocity over 24 hours. Superficial velocity was reduced to 0.3 mL/min after 24 hours of cell seeding phase. Cells were then cultured for two weeks in CM supplemented with 5 ng/mL FGF2. The culture medium was changed twice per week.

eVON was generated by seeding hAT-SVF cells at a ratio of 1:1 into pre-engineered eON after one week of osteogenic differentiation [42, 45]. After 24 hours of cell seeding phase, the superficial velocity was reduced to 0.3 mL/min for perfusion co-culture for additional two weeks in OM supplemented with 5 ng/mL FGF2. The culture medium was changed twice per week.

### Co-culture with MCF7 cells and fulvestrant treatment

MCF7-tdTomato/mVenus-p27K^-^ cells were seeded at 1:10 ratio onto eON, eVN, and eVON constructs under the perfusion bioreactor at 3 mL/min for 24 hours. Cells were co-cultured with the respective niches in CM at 0.3 mL/mmin for two weeks. Medium was refreshed twice weekly. Fulvestrant (100 nM) was added onto the respective niches after a week of co-culture, and refreshed twice weekly over a week.

### Cell isolation from the engineered niches (eON, eVN, and eVON)

The constructs were washed with phosphate-buffered saline (PBS) (Gibco; cat# 20012-027), and perfused at 3 mL/min superficial velocity with 0.3% collagenase type II (Worthington; cat# LS004176) in PBS for an hour at 37°C. Supernatants were collected and filtered through a 100 μm nylon mesh strainer (Corning; cat# 352360) and centrifuged at 500g for three minutes. Cells were then resuspended in FACS buffer (2% FBS (Gibco; cat# 10500-064), 2 mM ethylenediaminetetraacetic acid (EDTA) (Sigma; cat# E7889), PBS). Additional digestion with 0.05% trypsin-EDTA (Gibco; cat# 25300-054) under perfusion at 3 mL/min was performed for six minutes at 37°C. All fractions were pooled and, centrifuged at 500g for three minutes, and resuspended in FACS buffer.

### Flow cytometry

Phenotypic analysis of total MCF7 cells and mVenus-p27K^-^+ subpopulations was performed using CytoFLEX flow cytometer (Beckman Coulter) and BD FACSAria cell sorter (BD Biosciences). Cell suspensions were collected from the niche-free scaffold and the respective engineered niches. They were filtered through 30-µm mesh cell strainer tubes to obtain single cells. Debris exclusion and doublet discrimination were achieved by gating based on forward and side-scatter profiles and pulse width, respectively. Dead cells were identified and excluded using 4′,6-diamidino-2-phenylindole (DAPI) staining (DAPI+ cells). Cancer cells were determined by the positive expression of tdTomato. Single-stained and fluorescence minus one (FMO) controls were used to set gating parameters.

### Immunofluorescence staining

Medium was removed from the bioreactors, and the scaffolds with engineered niches were taken out of the bioreactors using tweezers. Scaffolds were rinsed with PBS, and fixed in 4% paraformaldehyde (PFA, Thermoscientific; cat# 28908) for 24 hours at 4°C. After the fixation, scaffolds were rinsed in PBS, and cut in half using a scalpel. One half of the scaffolds were used for whole-mount staining, and the other half were decalcified in 15% EDTA (0.5 M, pH: 7.2), renewed every other day for a week at 37°C with agitation. They were then embedded in paraffin, and sections were cut in 6µm thickness using Microtome (Thermo Scientific; cat# HM 340E). Tissue sections were hydrated in Ultraclear™ (J. T. Baker; cat# 3905.500PE), and a graded alcohol series. Slides were then subjected to heat-induced epitope retrieval (HIER) (pH: 6, Citrate Buffer, 1x, Quartett, AR-001-0120) for 15 min at 96°C. Sections were permeabilized and blocked with 3% bovine serum albumin (BSA) in PBST (0.2% Triton X-100 (Sigma; cat# 9002-93-1)) for an hour. Primary antibodies (*SI Appendix*, Table *S1*) were diluted in 0.5% BSA in PBST and incubated overnight at 4°C. Secondary antibodies (*SI Appendix*, Table *S2*) diluted in 0.5% BSA in PBST was applied for 30 minutes at room temperature. Slides were counterstained for a minute with DAPI (BD Biosciences; cat# 564907) solution. Slides were then mounted with Fluoromount™ aqueous mounting medium (Sigma; cat# F4680), and images were obtained using Nikon Ti2 microscope (NIS version 5.30.07) equipped with X-Light V3 confocal unit, photometrics Kinetix (29.4mm, back-illuminated sCMOS) camera and Apo Plan lambda 20x, NA0.75 objective. Images were stored and figures were prepared in OMERO [61] and image analyses were conducted using QuPath software (v0.5.1) [62].

### Whole-mount IF staining

Half of the fixed scaffolds were processed for whole-mount IF staining. Samples were permeabilized and blocked in 3% BSA (Sigma; cat# A9647), 0.2% Triton X-100 (Sigma; cat# 9002-93-1) in PBS for 6 hours at room temperature, followed by 72 hours of primary antibody incubation at 4°C with agitation. Antibodies were diluted in 0.5% BSA, 0.2% Triton X-100 in PBS and incubated overnight at 4°C. The following human primary antibodies were used: Anti-CD31 (Abcam; cat# ab9498), anti-NG2 (Abcam; cat# ab255811). After serial washes with PBST (0.2% Triton X-100 in PBS) and PBS, samples were incubated with secondary antibodies (Invitrogen) overnight at 4°C, washed again, and counterstained with DAPI. Samples were stored in PBS at 4°C until imaging. Confocal imaging was performed on a Nikon X-Light V3 spinning disk microscope using 0.9 µm z-step across ~150 µm total depth (~165 optical sections per scaffold). Image stacks were converted into maximum intensity projections using NIS-Elements software and further processed in OMERO software.

### Immunohistochemistry (IHC) staining of ERα

Formalin-fixed paraffin-embedded (FFPE) sections (6 µm) were stained for ERα (Thermo Scientific; cat# MA5-14501) using the Ventana Discovery Ultra (RocheDiagnostics) automated slide strainer. Briefly, tissue sections were deparaffinized and rehydrated, and were subjected to HIER, followed by incubation with the primary antibody (manually applied; 1 hour at 37°C). After washing, sections were incubated with the secondary antibody for 1 hour at 37°C. Detection was performed using the Ventana DISCOVERY ChromoMAP 3,3’-Diaminobenzidine (DAB) (Ventana, cat# 760-159) detection kit. Slides were then counterstained with hematoxylin II, followed by a bluing reagent (Ventana; cat# 790-2208, cat# 760-2037). The sections were dehydrated, cleared and mounted with permanent mounting medium. Slides were digitized using a Hamamatsu NanoZoomer S60 slide scanner equipped with a 40x objective (NA 0.95). Nuclear ERα+ MCF7 cell abundance (DAB+) was quantified by manual evaluation.

### Quantification of CD31+ endothelial network and NG2+ cell abundance

Maximum intensity projection images were generated using NIS-Elements software, subjected to ball-correction filtering to minimize signal to noise ratio. Files were then converted into pyramidal.ome.tiff format using ImageJ [63] software with Kheops plugin [64], and uploaded into QuPath. Full image annotation was created for each field of view (FOV). A single ANN pixel classifier to detect CD31+ (GFP channel) endothelial networks across each FOV was trained with examples of positive and negative signal with gaussian filter and sigma 2.0. Once the CD31+ endothelial network annotation was generated, a second pixel classifier was applied to detect NG2+ (Cy5 channel) structures within or adjacent to the CD31+ networks. The ratio of NG2+/CD31+ area per FOV was then calculated based on the workflow. The script containing the workflow was then run for all images, and the ratio of CD31+ endothelial network per FOV, and NG2+/CD31+ area per FOV was calculated based on the obtained measurements.

### Quantification of COL1A1, OCN deposition on IF staining images

IF staining images were converted into pyramidal.ome.tiff format using ImageJ software for compatibility with QuPath (v.0.5.1). Tissue boundaries were detected by the autofluorescence signal captured in the GFP channel (*SI Appendix*, Fig *S2B*) using a thresholder with sigma 5 and threshold 150. For COL1A1 quantification, a pixel classifier was created using a manual thresholding method on the Cy5 channel, where COL1A1 was visualized. Classifier resolution was set to full (0.56µm/px), with a Gaussian pre-filter applied and smoothing set to 0. The intensity threshold for COL1A1 and OCN positivity was empirically defined based on the signal from samples. Pixels above this threshold were classified as COL1A1+ ECM or OCN+ ECM, respectively. All classifiers were stored and batch-applied across full-section datasets.

### Quantification of tdTomato, mVenus, Ki67, and NR2F1 expression

IF staining images were first converted into pyramidal.ome.tiff format using ImageJ software, then uploaded into QuPath. Full image annotations were created to define FOVs. Nuclei were segmented using Watershed Cell Detection based on DAPI staining. Next, annotation for MCF7 cells (tdTomato+) were generated by creating a single measurement classifier based on the mean tdTomato signal intensity in the cytoplasm of MCF7 cells in the Cy3 channel. Thresholds were empirically defined through visual inspection. Following tdTomato+ cell annotation, cancer cells were segmented based on their nuclear expression of mVenus (GFP channel), Ki67 (Cy5 channel) and NR2F1 (Cy5 channel). Single measurement object classifier was created for each marker by applying channel-specific filters, and manually thresholding mean nuclear signal intensity to classify the positive cells. Cells co-expressing tdTomato with each nuclear marker were classified using composite object classifiers e.g., tdTomato+/Ki67+. Scripts for each marker panel were exported from QuPath and applied across all images. Manual curation was performed to remove staining artifacts and exclude false positives.

### Statistics

Data are presented as means ± standard deviation of the mean, and were analyzed by using GraphPad Prism software (v10.1.1). Unless otherwise stated/indicated, multiple (pairwise) comparisons were performed using Tukey’s test. Statistical significance was determined by one-way ANOVA followed by Tukey’s multiple comparison test. Statistically significant differences were defined as: *p<0.05, **p<0.01, ***p<0.001, ****p<0.0001.

## Supporting information

181225_Supplemental_Files_Bioarchive

## Acknowledgments

We thank past and present members of the Martin and Bentires-Alj laboratories for feedback and discussions. We also thank the Department of Biomedicine (DBM) Flow Cytometry Core Facility, specifically, Morgane Hilpert, Mihaela Barbu-Stevanovic, Jelena Markovic Djuric, Stella Stefanova for assistance with FACS. We are grateful to the DBM Microscopy Core Facility, particularly Loïc Sauteur, for assistance with imaging and image processing. We thank the DBM Histology Core Facility, particularly Diego Calabrese for performing the immunohistochemical staining. We thank T. Oki and T. Kitamura for providing the pMXs-IRES-puro/mVenus-p27K^-^ vector, and A. Bottos and N. E. Hynes for the pFU-Luc2-tdTomato vector. We also thank for past and present members of the Scherberich and Barbero laboratories for their helpful feedback and discussions. We are grateful to Noemi Torriero, Gangyu Zhang, and Benedetta Guagnini for their technical assistance, regular scientific exchange, and support. This work was supported by the European Commission under the Horizon Europea Marie Skłodowska-Curie Actions (MSCA) program (Grant No. 860715; SINERGIA), by the Freiwillige Akademische Gesellschaft Basel (FAG Basel), and by the Stiftung zur Förderung von chirurgischer Forschung und Spitalmanagement.

