## Supplementary material for "Bioengineered human bone-mimetic niche compositions regulate breast cancer cell quiescence and therapy response": 181225_Supplemental_Files_Bioarchive: 181225_Supplemental_Files_Bioarchive.pdf

[a]Laboratory of Tissue Engineering, Department of Biomedicine, University Hospital Basel, University of Basel, 4031 Basel, Switzerland; [b]Laboratory of Tumor Heterogeneity, Metastasis and Resistance, Department of Biomedicine, University Hospital Basel, University of Basel, 4031 Basel, Switzerland; [c]Microscopy Core Facility, Department of Biomedicine, University of Basel, 4031 Basel, Switzerland; [d]RESTORE Research Center, Université de Toulouse, INSERM 1301, CNRS 5070, EFS, ENVT, Toulouse, France.

[1]Equally contributing authors

*Ivan Martin and Mohamed Bentires-Alj

**Author Contributions:** E.C.K. conceived the study, designed and conducted experiments, analyzed and interpreted the results, and wrote the manuscript. M.G.M contributed to experimental planning, bioreactor experiments, data interpretation, and the manuscript refinement. S.L.F. contributed to experimental planning, data interpretation, and the manuscript refinement. E.M.B. developed and automated image quantification workflows. A.G.-G. contributed to data interpretation, and manuscript refinement. F.P. assisted with bioreactor experiments, histology, immunofluorescence staining, and imaging. A.D. contributed to histology and imaging. I.M. and M.B.-A. supervised the project. All authors read and provided feedback on the manuscript.

**Competing Interest Statement:** The authors declare no competing interests.

**This PDF file includes:**

Supporting text
Figures S1 to S5
Tables S1 to S2

**Other supporting materials for this manuscript include the following:**
Dataset S1 (separate file). QuPath scripts and classifiers used for the optimized IF staining image analysis pipelines

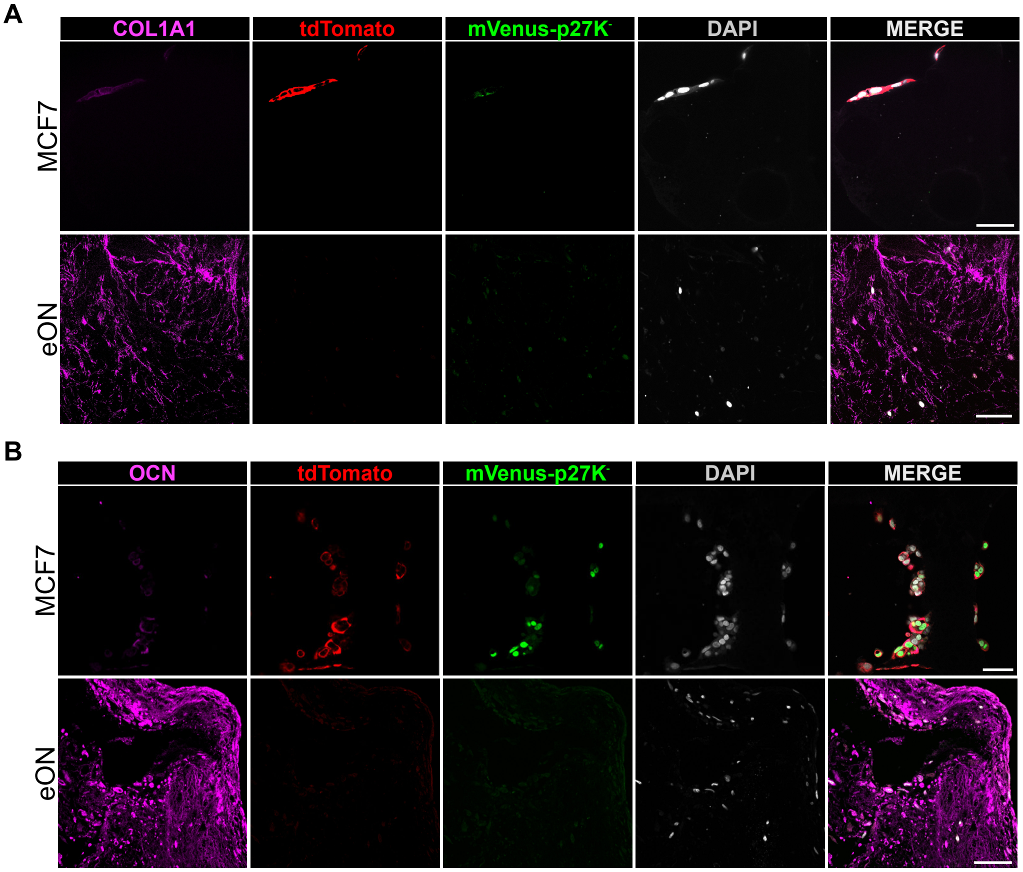


**Fig. S1. Osteogenic marker abundance confirms generation of the eON. (A, B)** Representative IF staining of **(A)** COL1A1 and **(B)** OCN in a niche-free scaffold and eON. Magnification: 20x; scale bar: 50 µm.

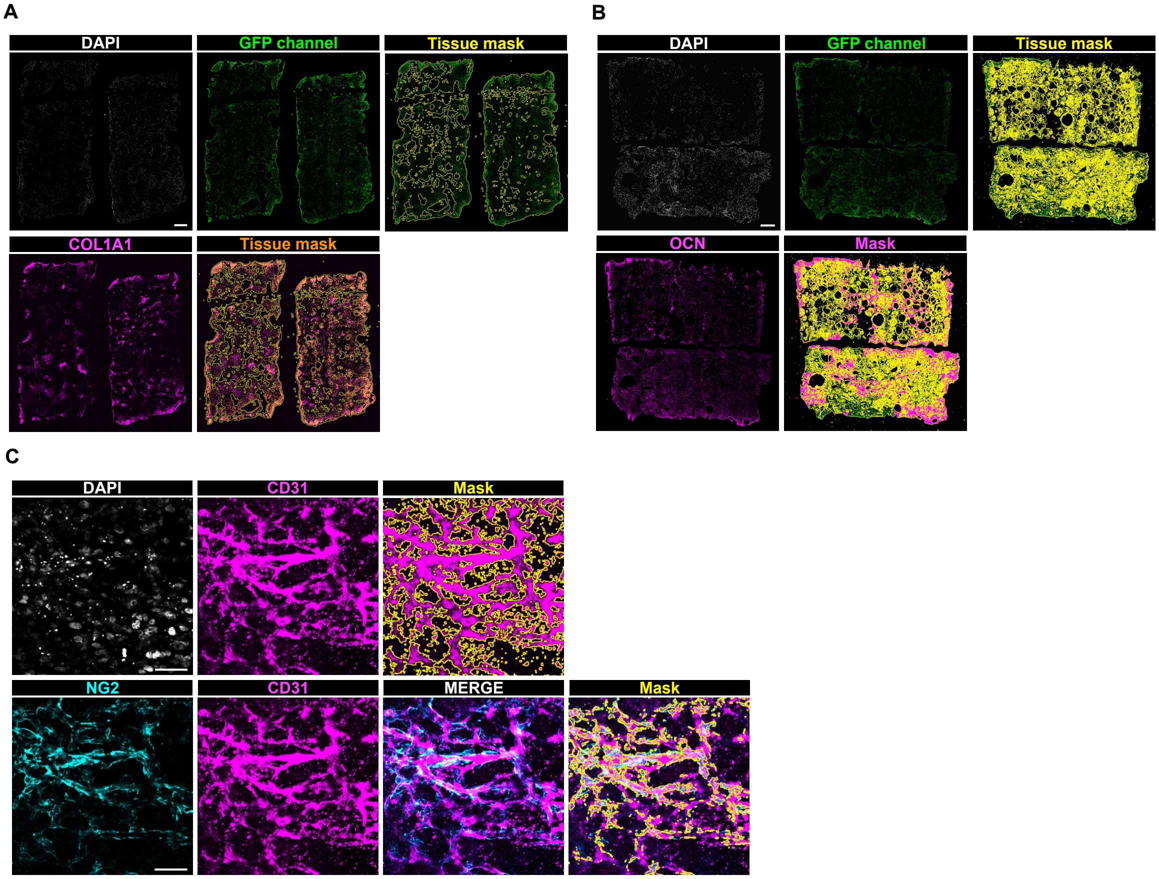


**Fig. S2. Optimized image analysis pipeline for quantification of osteogenic and vascular marker-positive area. (A)** COL1A1 (magenta) and tissue boundaries identified from autofluorescence in the GFP channel were used for pixel-based segmentation and quantification of ECM deposition using QuPath. **(B)** OCN (magenta) deposition within the co-culture scaffolds was similarly quantified using a manually thresholded pixel classifier. Masks indicate areas above threshold used for quantitative analysis. Images represent zoomed in FOVs from the respective niches after two weeks of co-culture with MCF7 cells. **(C)** Representative IF staining images and QuPath-based pixel classification for quantification of CD31+ endothelial networks (magenta) and NG2+ perivascular structures (cyan) from co-culture constructs. CD31+ endothelial network was used to segment vasculature-like structures, followed by overlay of NG2 signal. Masks indicate areas detected and quantified by classifier. Magnification: 20x; scale bar: 50 μm.

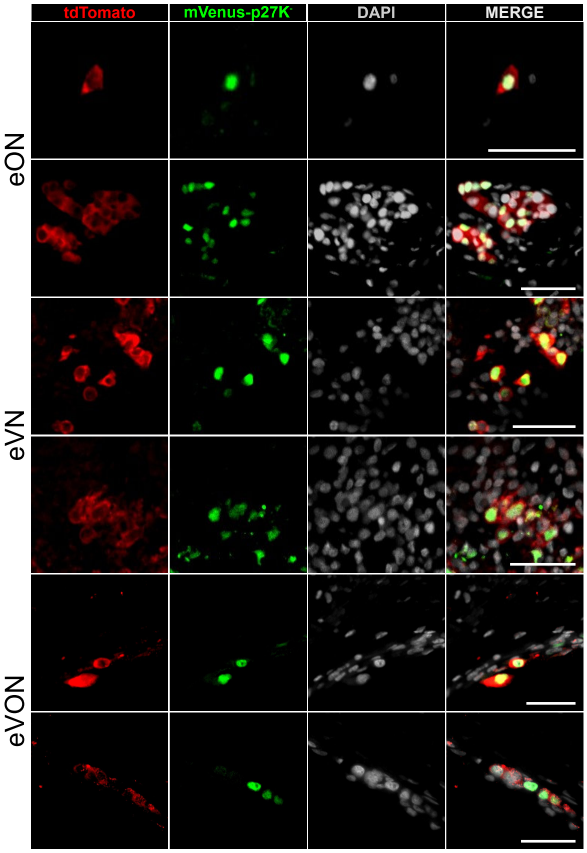


**Fig. S3. Quiescent cancer cells appear as solitary cells or as clusters in eON, eVN, and eVON.** Representative IF staining images showing quiescent MCF7 cells (tdTomato+/mVenus+; red and green) in the respective niches after two weeks of co-culture. DAPI marks nuclei (gray). Merged panels illustrate spatial distribution of quiescent cells across the engineered niches. Magnification: 20x; scale bar: 50 μm.

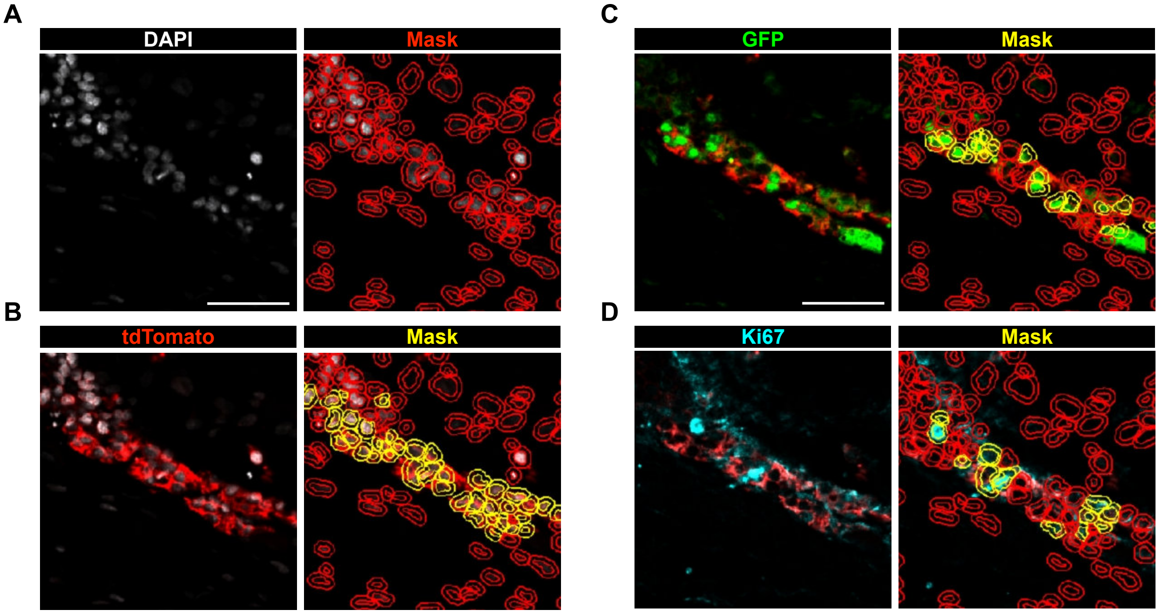


**Fig. S4. Optimized image analysis pipeline for quantification of quiescent and proliferative MCF7 fractions. (A-B)** QuPath-based cell detection (red mask) and cytoplasmic segmentation (yellow mask) used to quantify total number of cells and MCF7 cells (tdTomato+; Cy3 channel) per FOV. Quantification of **(C)** quiescent (mVenus+/Ki67-; GFP and Cy5 channel) MCF7 cells (yellow mask), and of **(D)** proliferative (mVenus-/Ki67+) MCF7 cells (yellow mask) by nuclear detection and intensity thresholding. Magnification: 20x; scale bar: 50 μm.

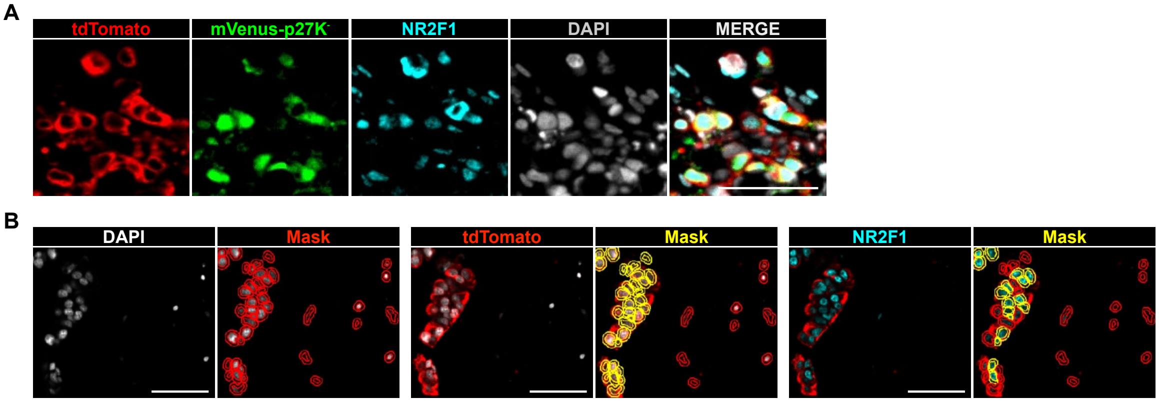


**Fig. S5. Nuclear colocalization of mVenus-p27K$^-$ and NR2F1 in MCF7 cells. (A)** Representative IF staining for tdTomato, mVenus, and NR2F1. **(B)** Optimized image analysis pipeline for quantification for nuclear NR2F1 abundance in MCF7 cells in the engineered constructs. QuPath-based cell detection (red mask) and cytoplasmic segmentation (yellow mask) used to quantify total number of cells and MCF7 cells (tdTomato+; Cy3 channel) per FOV. Quantification of nuclear NR2F1 expression (Cy5 channel) in MCF7 cells (Nuclear NR2F1+ CMF7 cells; yellow mask) by nuclear detection and intensity thresholding. Magnification: 20x; scale bar: 50 µm.

**Table S1. Primary antibodies used for IF staining**

| Antigen | Host species | Dilution | Supplier | Catalog Number |
|---|---|---|---|---|
| COL1A1 | Rabbit | 1:100 | Abcam | 138492 |
| OCN | Rabbit | 1:100 | Abcam | 93876 |
| RFP | Rabbit | 1:100 | Rockland | 600-401-379 |
| Cytokeratin (8/18) | Mouse | 1:200 | Novacastra | NCL-L-5D3 |
| GFP | Chicken | 1:1000 | Genetex | GTX13970 |
| Ki67 | Rat | 1:200 | Thermo Scientific | 14-5698-82 |
| COUP TF1 (NR2F1) | Rabbit | 1:100 | Abcam | ab181137 |

**Table S2. Secondary antibodies used for IF staining**

| Antibody | Host species | Dilution | Catalog Number | Supplier |
|---|---|---|---|---|
| Goat anti-Chicken IgY (H+L), Alexa Fluor 488 | Goat | 1:400 | A-11039 | Thermo Fisher |
| Goat anti-Mouse IgG (H+L), Cross-Adsorbed, Alexa Fluor 488 | Goat | 1:400 | A-11001 | Thermo Fisher |
| Goat anti-Mouse IgG (H+L), Highly Cross-Adsorbed, Alexa Fluor 555 | Goat | 1:400 | A-21424 | Thermo Fisher |
| Goat anti-Rabbit IgG (H+L), Highly Cross-Adsorbed, Alexa Fluor 555 | Goat | 1:400 | A-21429 | Thermo Fisher |
| Goat anti-Rat IgG (H+L), Cross-Adsorbed, Alexa Fluor 647 | Goat | 1:400 | A-21247 | Thermo Fisher |
| Goat anti-Rabbit IgG (H+L), Cross-Adsorbed, Alexa Fluor 647 | Goat | 1:400 | A-21244 | Thermo Fisher |

**Dataset S1 (separate file).** QuPath scripts and classifiers used for the optimized IF staining image analysis pipelines
